# CoLoRd: Compressing long reads

**DOI:** 10.1101/2021.07.17.452767

**Authors:** Marek Kokot, Adam Gudyś, Heng Li, Sebastian Deorowicz

## Abstract

The costs of maintaining exabytes of data produced by sequencing experiments every year has become a major issue in today’s genomics. In spite of the increasing popularity of the third generation sequencing, the existing algorithms for compressing long reads exhibit minor advantage over general purpose gzip. We present CoLoRd, an algorithm able to reduce 3^rd^ generation sequencing data by an order of magnitude without affecting the accuracy of downstream analyzes.

---

In the last few years, we have seen a rapid development of the third generation sequencing. The ability to produce very long reads makes Oxford Nanopore and PacBio technologies indispensable in genome assembly [6] or identification of long structural variants [18]. Maintaining gigantic collections of sequencing reads, which often reach hundreds of gigabytes per experiment, has become a major challenge and motivated intense research on compression techniques. Due to memory constraints, which prevented from maintaining the entire read collection in main memory, the early algorithms [7, 1, 16] were only slightly better than general-purpose compressors. The more recent methods [5, 17, 11, 2] successfully tackled this issue by clustering reads originating from close genome positions. All these approaches, though, were designed for short, high-quality Illumina reads and are unsuitable for reads from 3^rd^ generation instruments which are orders of magnitude longer and have different error profile. Among few algorithms able to compress ONT/PacBio FASTQ files, like SPRING [2], ENANO [3], or LFQC [13], none takes advantage of the data redundancy in the overlapping reads. This, together with the lossless compression of the quality stream, which is built over significantly greater alphabet than DNA (94 instead of 4 symbols) and exhibits larger noise, limits advantages over, de facto standard, gzip on 3^rd^ generation data to tens of percent.

In the paper, we present CoLoRd, a compression algorithm for ONT and PacBio sequencing data. Its main contributions are (i) novel method for compressing the DNA component of FASTQ files and (ii) lossy processing of the quality stream. The idea of the former is based on the overlap graphs [12] often employed by long-read assemblers [9, 8]. As CoLoRd does not aim at finding true overlaps (which are critical for genome reconstruction), but those convenient for differential compression of reads, the throughput of the overlap detection can be increased at the cost of the accuracy. The algorithm determines *k*-mer similarities between the reads and use them for identifying closest neighbours — potential references in the differential compression. The compression itself is based on *anchors*, i.e., exact sequence matches. The between-anchor areas are further scanned for similar fragments. As a result, the algorithm translates a read to an edit script describing differences w.r.t. similar reads (Figure 1a), which is then entropy-encoded. The presence of priority modes, i.e., *memory* (default), *balanced*, and *ratio* which differ mainly by the fraction of reads used as references, allows suiting the algorithm to particular needs.

**Figure 1:**
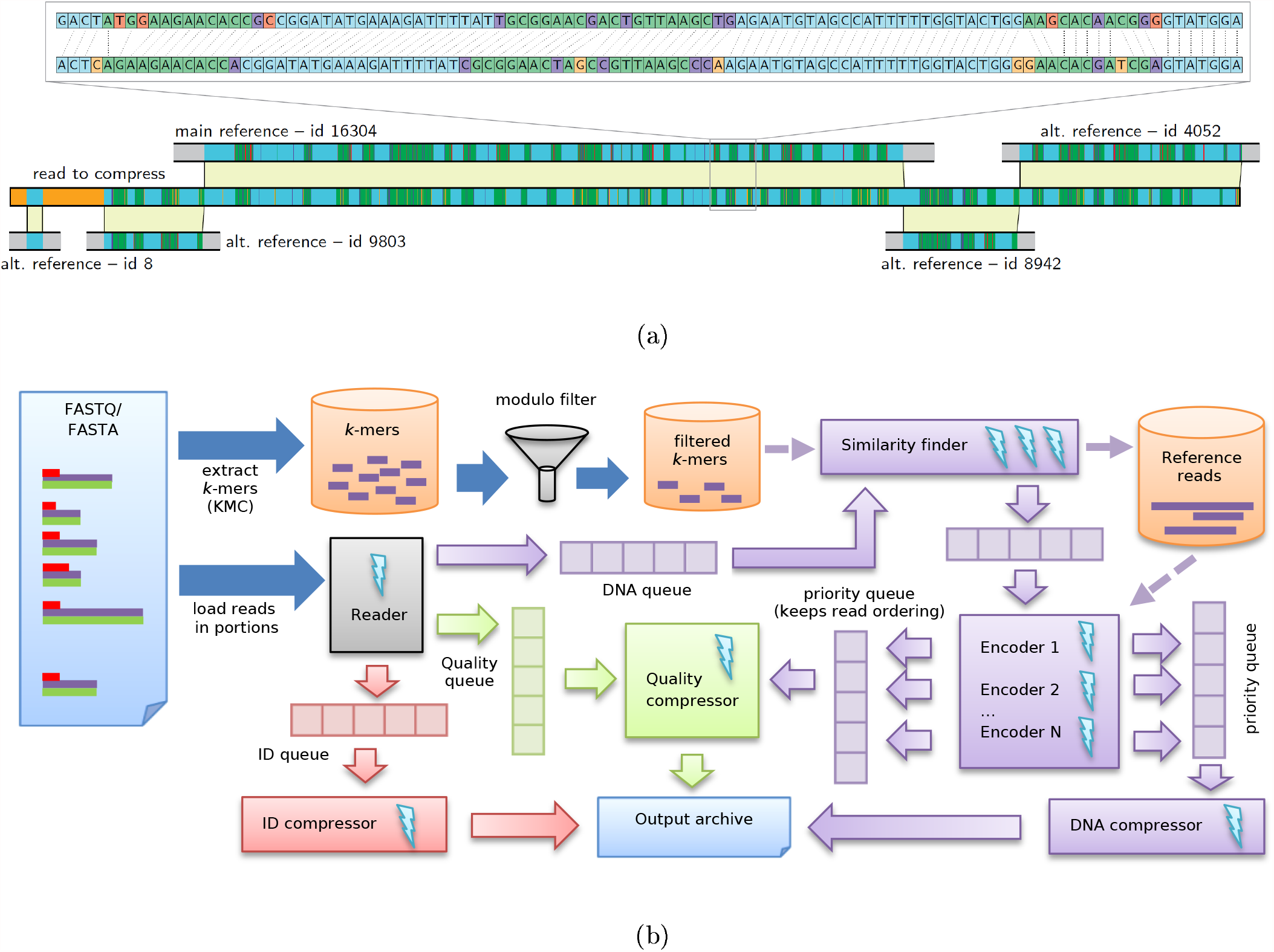
CoLoRd operation principles. **a** Encoding a query read w.r.t. to the reference reads. The edit script was generated while compressing a read from Zymo dataset. The colours indicate anchors (blue), matches (green), mismatches (purple), insertions (orange), deletions (red), and unmatched areas (gray). **b** Processing scheme of DNA (purple), quality (green), and identifiers (red) streams. Lightnings represent computing threads.

The encoding of the quality stream is done with a use of the context-based approach inspired by ENANO, complemented with the knowledge of read overlaps. In agreement with the findings on Illumina data [17], we show that the resolution of quality scores can be reduced without affecting the accuracy of downstream analyses. Therefore, quality levels are by default subject to the lossy compression. CoLoRd uses 4-level quality binning for Nanopore data with a scheme of [Q0, Q6], [Q7, Q13], [Q14, Q25] and [Q26, Q93]. For HiFi data, Q93 value is stored in a separate bin. CoLoRd calculates a representative quality in each bin such that the average read quality is preserved. The lossless mode, together with other variants, are also provided. The functionality of our algorithm is complemented by the presence of reference-based compression, which further improves ratios. CoLoRd was implemented in the C++ programming language and takes advantage of multiple computing threads (Figure 1b).

The advantage of CoLoRd over general-purpose (gzip, 7zip) and specialized (ENANO, SPRING) compressors was confirmed on 24 read sets representing different technologies (ONT, PacBio) with sizes ranging from 2 to 786 GB (Methods). As presented in Figure 2a, our algorithm revealed its potential on the latest ONT Bonito-base-called and PacBio HiFi data sets. By efficiently finding overlaps in the high fidelity reads, CoLoRd reduced the size of the DNA stream by two orders of magnitude. This, accompanied by the quantization of quality levels, allowed squeezing FASTQ files to the 1/25 of their original size, which translated to 4-fold (ONT Bonito) and 10-fold (HiFi) advantage over lossless gzip compression. For instance, HiFi HG002 30 *×* data set was compressed from 219 to 10 gigabytes. With further accuracy improvements in 3^rd^ generation sequencing on the horizon, even better results could be achieved. Contrarily, the high error rate of Nanopore reads produced by the older base callers, bounded the advantage of CoLoRd to lower, though, still significant levels. For experimental integrity, we additionally simulated lossy compression in gzip, 7zip, ENANO, and SPRING by binning quality levels in the input files. In this scheme, our algorithm was ahead of the rest with 2–4-fold improvement over gzip. In the reference-based mode, CoLoRd offered even better compression ratios and was superior to other algorithms using reference genomes like RENANO [4] or CRAM [1] (Supplementary Table 1).

**Figure 2:**
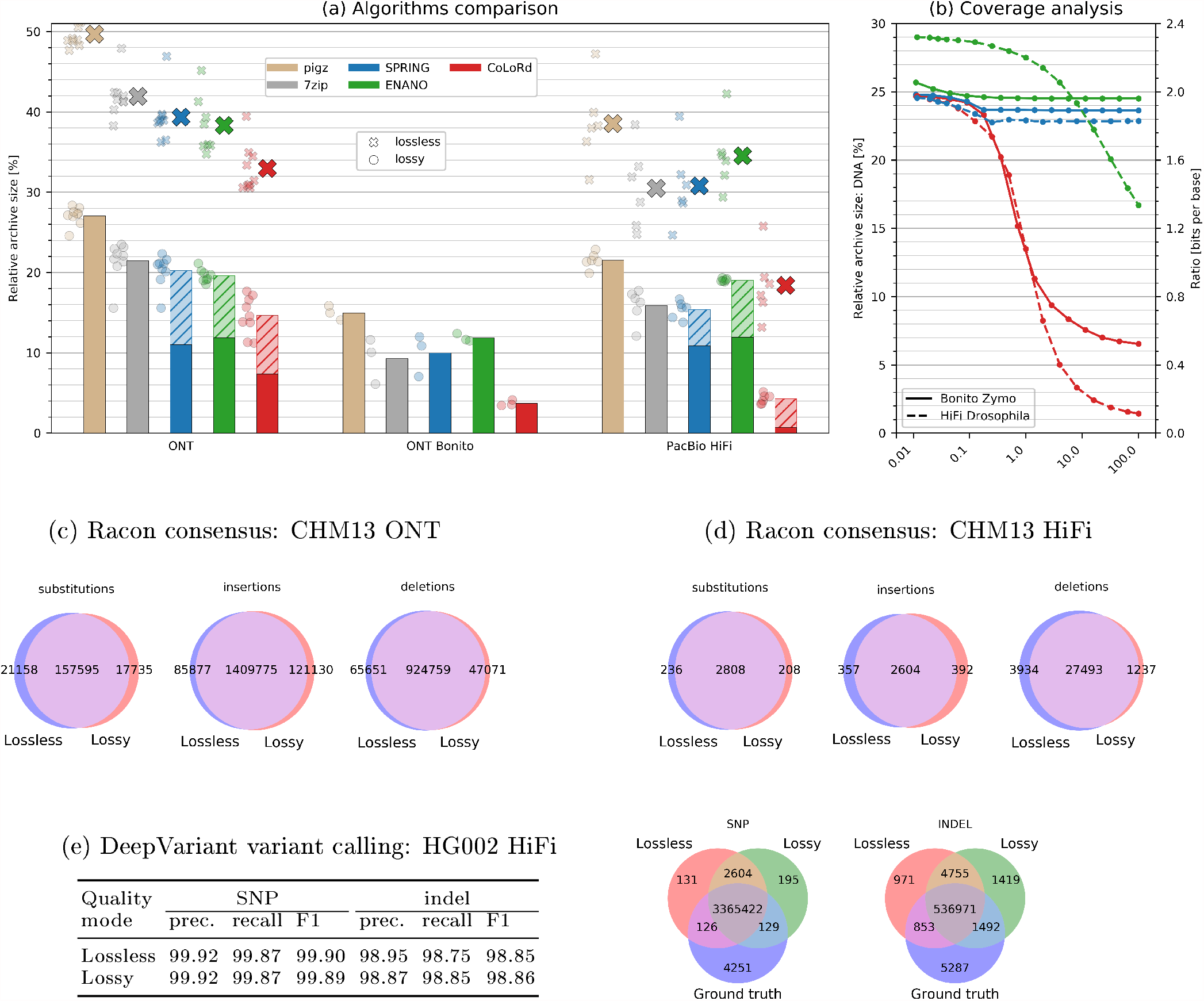
Analysis of lossless and lossy compression. **a** Average performance of the algorithms with lossless (large crosses) and lossy (bars) quality compression (hatched areas represent contribution of the quality stream). The results on the individual data sets are presented as small crosses (lossless) and circles (lossy). The lossy archives for pigz, 7zip, SPRING, and ENANO were generated by binning quality scores in the input files. ENANO failed to process the largest of the ONT datasets. **b** Relative size of the DNA archive as a function of sequencing coverage evaluated on the selected ONT and HiFi data sets. The corresponding compression ratio in bits per base is presented on the right vertical axis. **c, d** Results of the Racon consensus generation expressed by the number of substitutions, insertions, and deletions w.r.t. reference genome. **e** Accuracy of variant calling with DeepVariant.

An essential property of CoLoRd is that with increasing coverage, the number of overlaps, thus the efficiency of differential compression, grows as well. As the analysis of two selected data sets shows (Figure 2b), when taking 0.01% of reads, all investigated algorithms compressed the DNA stream to approximately 1/4 of its initial size. For the entire read set, however, CoLoRd offered 20-fold (Bonito Zymo with Q20 chemistry) and 100-fold (HiFi Drosophila) reduction of the DNA component, while the competitors exhibited minor or no improvements in the compression ratio.

Importantly, superior compression results of the presented algorithm came at competitive running times. The lossy compression/decompression speeds at 24 threads were way above 50 MB/s, which is the throughput of PromethION—the fastest among 3^rd^ generation instruments. Memory requirements were also reasonable with 16 GB for the aforementioned HG002 data set.

Detailed results including all CoLoRd priority modes and CLR data sets are presented in Supplementary Worksheet.

We evaluated consensus and variant calling accuracy to see how much lossy quality compression may affect downstream analyses. We mapped CHM13 HiFi reads to the CHM13 telomere-to-telomere (T2T) assembly v1.1 [14], generated the consensus with Racon v1.4.20 [19], aligned the consensus to the assembly with minimap2 v2.20 [10] and counted consensus differences (Figure 2c-d). The T2T assembly was derived from multiple data types including HiFi, Nanopore and Illumina reads. Our HiFi-only Racon consensus from raw reads differs from the T2T assembly at a rate of one difference per 1kb. Interestingly, lossy quality compression slightly improves the consensus accuracy to one difference per kb, suggesting Racon might be more compatible with our lossy scheme. We also applied the same procedure to Nanopore reads and observed that lossy quality compression does not reduce the Racon consensus accuracy, either (one difference per 1.1kb for both lossless and lossy compression). To evaluate variant calling accuracy, we mapped HG002 HiFi reads to the human reference genome GRCh8 and called small variants with DeepVariant v1.1.0 [15]. When compared against the Genome-In-A-Bottle v4.2.1 truth [20], variant calls from lossy quality compression are as accurate as calls from raw reads (Figure 2e). With these experiments, also given the fact that most long-read assemblers ignore base quality at the assembly step, we conclude that lossy quality compression does not affect the accuracy of most downstream analyses. Detailed investigation of different quality compression modes is presented in Supplementary Table 2.

Here, we introduce CoLoRd, a comprehensive package for compressing Oxford Nanopore/PacBio sequencing data. Equipped with an overlap-based algorithm for compressing the DNA stream and a lossy processing of the quality information, it allows even tenfold space reduction compared to gzip, without affecting down-stream analyses like variant calling or consensus generation. This opens new opportunities in the field of long read sequencing, where maintaining gigantic data volumes has become one of the major contributor to the overall costs.

## Supporting information

Supplementary Worksheet

## Acknowledgements

The work was supported by National Science Centre, Poland, project DEC-2019/33/B/ST6/02040 (MK, AG, SD) and by US National Institutes of Health (grant R01HG010040, U01HG010971 and U41HG010972 to HL).

## Author information

MK and SD designed and implemented the algorithm. HL designed the variant calling/consensus analyzes, investigated and described their results. AG, SD, and MK conducted the experiments. AG prepared visualizations and wrote the majority of the manuscript. All authors read and approved the final manuscript.

## Competing interests

HL is a consultant of Integrated DNA Technologies, Inc and on the Scientific Advisory Boards of Sentieon, Inc and Innozeen Inc.

## Methods

### CoLoRd overview

CoLoRd compressor handles reads in FASTA or FASTQ format. In the former case, the input consists of two independently compressed streams: DNA data, which contribute predominantly to the file size, and sequence headers. In FASTQ files there is an additional base quality stream of the same size as DNA data.

The main contribution of the CoLoRd is the DNA compression algorithm. DNA sequences of reads are built upon 5-element alphabet (A, C, G, T, N) and are highly redundant due to: (i) same genome fragments being covered by multiple reads and (ii) similarity between different genome regions. CoLoRd is the first method which takes advantage of the characteristics of Oxford Nanopore and PacBio data when compressing DNA stream. The algorithm processes reads one by one determining their *k*-mer similarity to already analyzed reads. The resulting similarity graph is then used to identify closest neighbours which are references for differential compression of the current read. The procedure is based on *anchors*—exact sequence matches determined with a use of *m*-mers (*m ≤ k*). The between-anchor areas are further scanned for similar fragments. As a result, the algorithm translates a read to an edit script describing differences w.r.t. similar reads. Finally, the script is entropy-encoded. This multi-step approach renders superior compression ratios of DNA data.

Quality stream consumes same number of bytes in FASTQ file as DNA data (there is one quality symbol per base). However, base quality in 3^rd^ generation sequencing is expressed by one of 94 values and is non-redundant, by nature. Therefore, the quality data can be compressed losslessly to much less extent than DNA. CoLoRd employs a context-based approach inspired by ENANO [3] for this purpose. However, in many analyses base quality can be discretized to fewer values or even discarded at all. In particular, the common practice is to threshold scores at assumed level distinguishing only between “high” and “low” quality. Therefore, a lossy compression mode for quality was introduced in CoLoRd allowing significant reduction of this stream.

Since sequence headers contribute marginally to the sizes of FASTA/FASTQ files, they are compressed with well-established token-based method analogously as in FQSqueezer [23]or ENANO.

In the following subsections a detailed description of selected compression aspects is presented.

### Similarity graph construction

The initial algorithm step is filtering *k*-mers of the input reads. For this purpose KMC package [28] is executed with *k* automatically adjusted to the data set. *K*-mers with less than *L* = 4^∗^ (possibly, sequencing errors), or more than 80 occurrences (non-informative) in the entire read set are discarded. Finally, to reduce the representation, we select subset *U* of *k*-mers, such that *U* = {*u* : *h*(*u*) mod *f* = 0} with *h* being a Murmur3 hash function and *f* equals to 12 (ONT) or 40 (HiFi). Parameter *f* balances the sensitivity and the memory footprint of the procedure.

As a next step, a graph *G* is constructed with reads being the vertices and edges representing *k*-mer similarities. A weight of an edge is a number of common *k*-mers in reads representation. An edge is directed from a read analyzed later towards one analyzed earlier. For memory efficiency, each vertex has at most *max*_*candidates* = 8 outgoing edges to the most similar reads.

The graph is built with a use of *kmers*2*reads* association table which maps *k*-mers to lists of read identifiers they are contained in. The structure is initially empty. The algorithm processes reads one by one according to the following procedure:

1. Extract all unique *k*-mers from a current read *r*.
2. Create an association table *M* indexed by read identifiers to store number of common *k*-mers between *r* and other reads.
3. For each *k*-mer *u* ∈ *r* ∩ *U*:
  a. Get list *L*_*id*_ of read identifiers containing *u* (if any): *L*_*id*_ ← *kmers*2*reads*[*u*].
  b. For each identifier *id* ∈ *L*_*id*_ update *M* table: *M* [*id*] ← *M* [*id*] + 1
  c. Indicate in *kmers*2*reads* that *u* was contained in *r*: *L*_*id*_ ← (*L*_*id*_, id(*r*)).
4. Sort table *M* in a descending order w.r.t. the number of common *k*-mers.
5. Add to the graph *G* a new vertex representing read *r*.
6. Add directed edges from the new vertex to *max*_*candidates* reads with most *k*-kmers in common (use the number of *k*-mers as a weight).

### Anchor-based DNA encoding

The compression algorithm checks if consecutive reads contain neighbours in the similarity graph *G*. If not, a current read *r* is compressed in a plain mode. Otherwise, a differential compression between the read and the neighbours is performed with a use of *anchors*, i.e., exact matches between sequences. The anchors are determined as follows:

1. Establish all *m*-mers of *r* (*m* is automatically adjusted to the data set).
2. If *r* contains less then |*r*| */*2 unique *m*-mers (|*r*| being the read length), consider it as highly repetitive and encode in the plain mode.
3. Otherwise, determine *m*-mers of all reads in the neighbourhood of *r* in the graph *G*.
4. Establish intersections of *m*-mers from *r* and each of its neighbours. An intersection is represented as a list of common *m*-mers and two lists of their positions in *r* and its neighbour *s*. The reverse complements of the neighbours are also investigated.
5. Using common *m*-mers as alphabet symbols, determine the longest common subsequence (LCS) of *r* and *s*.
6. Join overlapping *m*-mers from LCS to obtain anchors between *r* and *s*.

As a next step, *r* is encoded with respect to its neighbour *s*_best_ with the largest sum of anchor lengths. The algorithm alternately process non-anchor and anchor fragments of *r* and *s*_best_. Anchors are encoded as lengths and positions in *s*_best_. For non-anchor fragments, an edit script is determined with a use of Edlib library [31]. Then, based on the entropy estimation, the decision is made whether the edit script is used for representing the analyzed read fragment. If not, two scenarios are possible. (i) If the fragment is at least 64 symbols long, it is treated as a temporary read *r′* and an attempt is made to encode it with respect to an alternative candidate 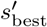 with the largest total anchor length w.r.t. *r′*. (ii) If the fragment is shorter than 64 or no alternative candidates for *r′* have been found, it is encoded in the plain mode. Up to 3 levels of recurrence are allowed by the procedure.

### Entropy encoding of DNA data

The edit script from the previous step is a sequence of the tuples of the types: *start_edit_script, start_plain, start_plain_with_Ns, anchor, match, insertion, deletion, substitution, skip, alternative_ref, main_ref, plain, end_of_read*. The first three of them can start the edit script and we entropy encode the starting tuple type taking as a context 4 previous read-starting-tuple types.

When we notice *start_plain* or *start_plain_with_Ns*, we store also the read length. Then, we have to encode the symbols A, C, G, T, or additionally N (in the later case). We store them using an entropy coder taking as a context (in case of plain read) or 4 (in case of plain read with Ns) previous symbols.

The processing of truly-edit-script-described reads is more complex. First of all we store the main reference read id and information whether the reference read is taken as it is or it is reverse complemented. Then, for each successive tuple we entropy encode its type. The context is formed of 3 previous tuple types, 2 preceding symbols in the current read and the information whether recently we have (w.r.t. reference read): medium-size insertion (from 2 to 100 symbols), long insertion, medium-size deletion, long deletion, nothing of above. The remaining of the tuple is encoded depending of the type:

- *alternative_ref*— it can happen that the same alternative reference read id will appear more than once in the edit script for the current read. Therefore, first we encode the information whether this read id has already been seen in the current edit script. If so, we store just its short “local” id (small number). Otherwise, we store its read id and the information whether it should be reverse complemented (just to serve as a reference for the current read) or not,
- *anchor*— the anchor length is entropy encoded,
- *match* — nothing needs to be encoded here,
- *insertion* — the inserted symbol is entropy encoded. The context is formed of: the symbol at the corresponding position in the reference read and 4 previous symbols,
- *deletion* — nothing needs to be encoded here since this tuple denotes just a single-symbol deletion w.r.t. a reference read,
- *substitution* — the symbol is entropy encoded. The context is formed of: corresponding symbol in the reference read, 3 previous symbols in the current read. Additionally, in the *ratio* priority, the context is extended by an information whether the 3rd and 4th previous symbols in the current read are same,
- *skip* — this tuple type denotes that the current position in the reference read should be changed by more than 16 positions. We entropy encode the skip length,
- *main_ref*— nothing needs to be encoded here since this tuple type denotes that switch from an alternative reference read to the main reference read,
- *end_of_read*—nothing needs to be encoded.

### Quality scores

#### Lossless compression

In the lossless mode, the qualities are compressed as they are. The context is formed of several parts. First, we take two previous quantized quality values. The quantization is made to reduce the alphabet from 94 possible values to 11 values. Then, we extend the context by 4 DNA symbols from the read from: the corresponding position, two previous positions, following position. This strategy is similar to the one used in ENANO. We, however, do not gather similar contexts. Moreover, we use information from the analysis of the DNA part of the read. In *balanced* and *ratio* modes we extend the context by an information, whether the DNA symbol was encoded as: a match, an anchor, something else.

#### Lossy compression

The lossy quality compression must take into consideration two major concerns. The first one is the ability to reconstruct with some resolution the information about quality of individual base calls, which may be useful in the applications like variant calling. At the same time, it is important to preserve as accurately as possible the average read quality in a case it is employed for read filtering. As a result, it was decided to map original levels into bins representing *poor, moderate, good* and *very good* bases. The default boundaries of the bins and values representing them were established experimentally as [Q0, Q6] → Q3, [Q7,Q13] → Q10, [Q14,Q25] → Q18, [Q26,Q93] → Q35 (for HiFi data, Q93 value is stored separately due to its special meaning in some applications). The context is formed of 6 previous, quantized, quality values. It is also extended by 4 DNA symbols and information about encoding, similarly as in the lossless processing. The aforementioned lossy compression modes are referred to as *4-fixed* (ONT) and *5-fixed* (HiFi).

In Figure 3 we present on selected data sets how the binning affects average read quality. In the case of Zymo2 and Zymo2 R10 samples, the average read qualities were upper-bounded by Q15, thus the error introduced by binning was small on the entire range of reported Q levels. In particular, the average read qualities were mostly underestimated by less then 0.75 (Zymo2) and 0.5 (Zymo2 R10). The different situation was in the case of lun and NA12878 data sets, where the range of Q was larger (up to Q35 and Q50, respectively). The underestimation of average quality clearly increased with rising Q, which was expected as the wide range of high quality calls [Q26,Q93] is represented by a single value Q35. Importantly, at the level Q20, which is often considered as a threshold for *good* calls (1% error probability), the average quality was underestimated slightly (between 0.5 and 3 in the vast majority of reads). This, however, may still be unacceptable when read filtering based on the average quality is applied. Therefore, an additional *4-avg* /*5-avg* modes were proposed in which value representing a bin is the average of all read base calls falling into that bin stored. During decompression, a floor and ceiling of this value are outputted in the appropriate proportion in order to accurately reconstruct the average read quality. As this feature renders only slightly larger archives compared to the fixed bin representation, it is used by default in CoLoRd.

**Figure 3:**
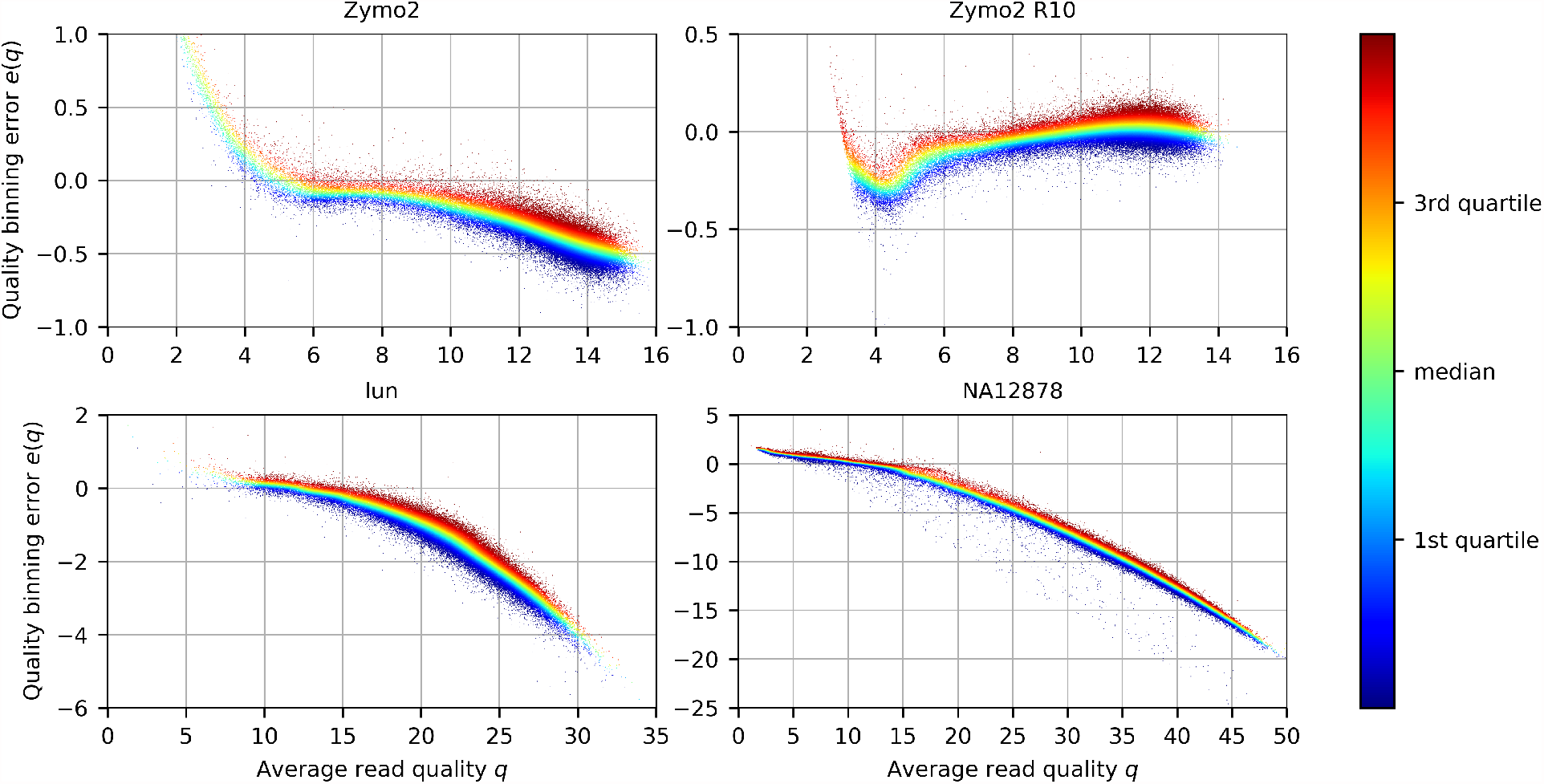
Distribution of read quality errors *e*(*q*) after binning w.r.t. average read quality *q*. Each read is represented as a point. The color of a point (*q, e*(*q*)) is calculated by taking all points from [*q −* 0.5, *q* + 0.5] neighbourhood, sorting them w.r.t. *e*(*q*), and taking its relative position in the resulting sequence.

To understand how *4-avg* mode works, let us assume that the quality part of a read is:

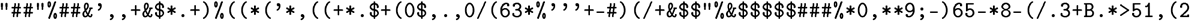

In compression, we calculate averages of symbols in ranges [Q0, Q6], [Q7, Q13], [Q14, Q25], [Q26, Q93]. Let us denote them as *a*_7_, *a*_14_, *a*_26_, *a*_93_. They are stored with precision 1*/*256. During decompression we load the stored values, which for the given example are: *a*_7_ *≈* 3.3984375, *a*_14_ *≈* 9.6484375, *a*_26_ *≈* 17.8125, *a*_93_ *≈* 29.33203125. Then we decode bin numbers one by one are output quality score ⌊*a*_*x*_⌋ or ⌈*a*_*x*_⌉ to minimize the difference between the already decoded quality scores for a bin and the stored average score. In this way we obtain the scores:

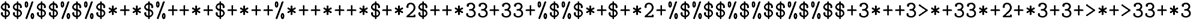

The additional quality modes in CoLoRd are:

1. *2 -fixed* : [Q0, Q6] → Q1, [Q7, Q93] → Q13,
2. *2-avg* : as above, but with averages as bin representatives,
3. *none* : all Q values mapped to Q0,
4. *avg* : all Q values mapped to the average read quality.

The bin boundaries and representatives in all modes can be redefined by the user according to his needs.

### Reducing execution time and memory requirements

By default, all reads without N symbols are used as potential references. This requires storing them in the memory which, in spite of packing four symbols per byte, results in excessive memory requirements for large sets of reads (e.g., human genome sequencing experiments). Therefore, we introduced a mechanism which decreases the probability of using a read as a potential reference as the compression proceeds. Let *R* indicate the reference genome length (which is estimated from *k*-mers). The idea is that after processing *gR*, 2*gR*, 3*gR*, … bases of the input file, with *g* being some parameter, the probability of adding current read to the collection of potential compression references decreases as 1, 1*/*2, 1*/*3, …, The memory saving concerns also *kmers*2*reads* structure, which does not have to store non-reference reads.

In order to maximize the utilization of available computational resources, CoLoRd is multi-threaded. The algorithm workflow is divided into independent blocks which exchange data using synchronized queues. In order to minimize synchronization overheads, the reads are processed in larger packs. The core of the processing consists of a thread pool whose elements, depending on the current workload, perform (i) similarity graph construction and (ii) anchor-based compression. Additionally, there are four threads responsible for entropy encoding of DNA, quality, and header streams as well as loading reads from FASTA/FASTQ files.

### Reference-based compression

The reference-based compression is based on the pseudo-reads generated by sliding the window over the reference genome with some overlap. The length of the window is 20 times greater than the average read length estimated during *k*-mers counting. The pseudo-reads are used as an initial set of reference reads in the previously described differential compression. By default, the reference genome is stored in the archive, making it self-contained. Alternatively, the genome file has to be specified explicitly during the decompression. In this case, an MD signature of the reference is stored in the archive to make sure that the proper reference is given.

### Datsets

The datasets used in the experiments are characterized in Table 1.

**Table 1:**
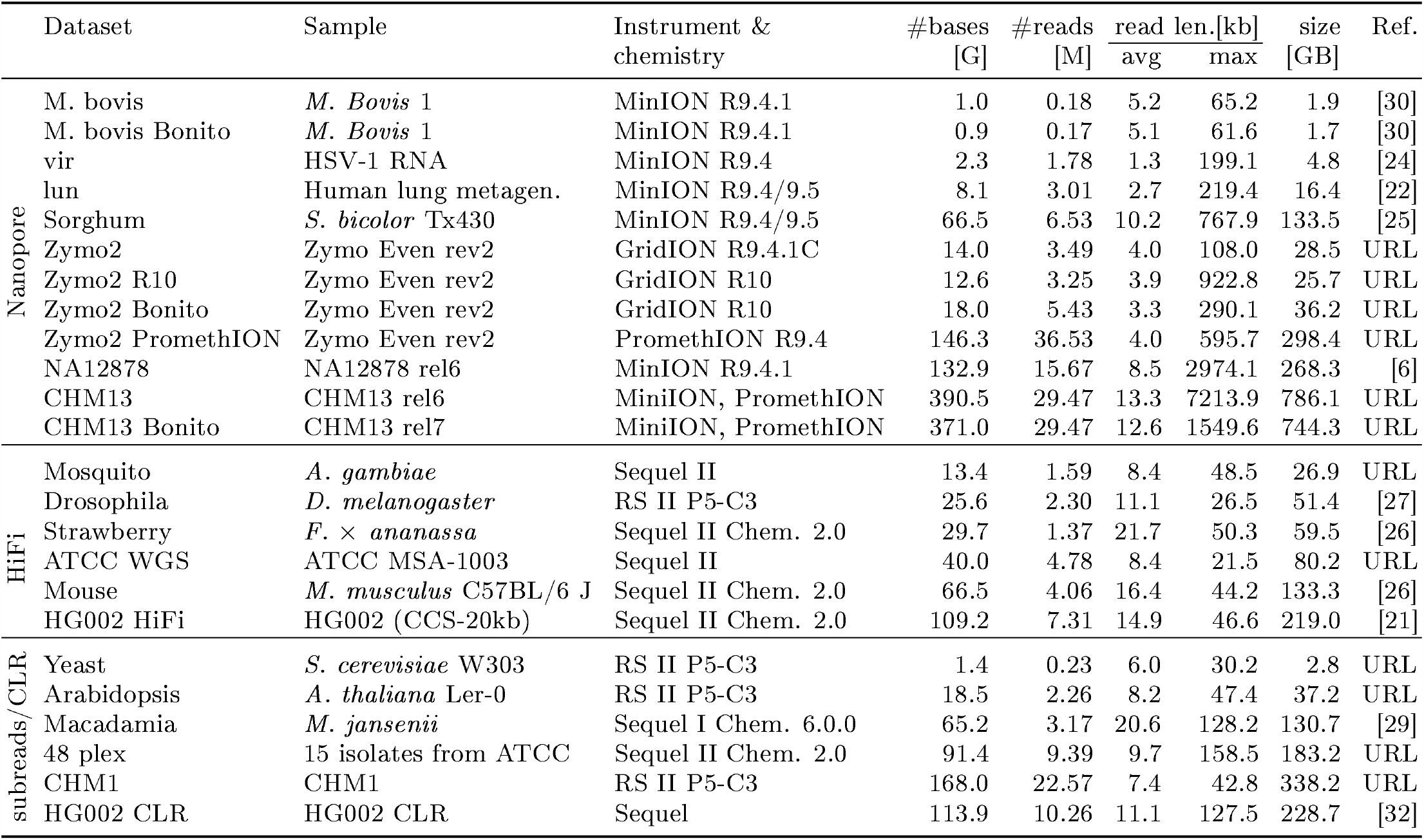
Characteristics of the data sets used in the study.

## Data availability

Oxford Nanopore data sets:

M. bovis: ERR4179765; M. bovis Bonito: ERR4179766; vir: ERR2708427-ERR2708436; lun: PRJEB30781; Sorghum: SRX4104135-SRX4104138; Zymo2: https://nanopore.s3.climb.ac.uk/Zymo-GridION-EVEN-BB-SN.fq.gz; Zymo2 R10 https://s3.climb.ac.uk/nanopore/Zymo-GridION-EVEN-BB-SN-PCR-R10HC-flipflop.fq.gz; Zymo2 Bonito ERR5396170; Zymo2 PromethION https://nanopore.s3.climb.ac.uk/Zymo-PromethION-EVEN-BB-SN.fq.gz; NA12878 : http://s3.amazonaws.com/nanopore-human-wgs/rel6/rel_6.fastq.gz; CHM13: https://s3-us-west-2.amazonaws.com/human-pangenomics/T2T/CHM13/nanopore/rel6/rel6.fastq.gz; CHM13 Bonito: https://s3-us-west-2.amazonaws.com/human-pangenomics/T2T/CHM13/nanopore/rel7/rel7.fastq.gz

PacBio HiFi datasets:

Mosquito: SRX8642991, SRX8642992; Drosophila SRX499318; Strawberry: SRR11606867; ATCC WGS: PRJNA546278; Mouse SRR11606870; HG002 HiFi: SRR10382244, [SRR10382245](https://www.ncbi.nlm.nih.gov/sra/SRR10382244), [SRR10382248](https://www.ncbi.nlm.nih.gov/sra/SRR10382244), [SRR10382249](https://www.ncbi.nlm.nih.gov/sra/SRR10382244)

PacBio CLR datasets:

Yeast: http://gembox.cbcb.umd.edu/mhap/raw/yeast_filtered.fastq.gz;

Arabidopsis: http://gembox.cbcb.umd.edu/mhap/raw/athal_filtered.fastq.gz; Macadamia: SRR11191909;

48 plex: https://downloads.pacbcloud.com/public/dataset/MicrobialMultiplexing_48plex/48-plex_sequences/;

CHM1: http://datasets.pacb.com/2014/Human54x/fastq.html; HG002 CLR: SRX5590586

Other:

CHM13 T2T assembly v1.1: GCA_009914755.3;

GRCh38 assembly: GCA_000001405.15_GRCh38_no_alt_analysis_set.fna.gz; Genome-In-A-Bottle v4.2.1

HG002 reference variants:

https://ftp-trace.ncbi.nlm.nih.gov/ReferenceSamples/giab/release/AshkenazimTrio/HG002_NA24385_son/NISTv4.2.1/GR

## Code availability

The source code of CoLoRd is available at https://github.com/refresh-bio/CoLoRd.

## Footnotes

∗ The parameters differ across priority modes (*memory*/*balanced* /*ratio*) and technologies (0NT/HiFi/CLR). For simplicity, we give the values in a default priority mode (*memory*) for 0NT and, optionally, HiFi, when it differs significantly.

## Notes

https://github.com/refresh-bio/CoLoRd

### Competing Interest Statement

Heng Li is a consultant of Integrated DNA Technologies, Inc and on the Scientific Advisory Boards of Sentieon, Inc and Innozeen Inc.

