## Supplementary material for "CoLoRd: Compressing long reads"

July 17, 2021

1. Faculty of Automatic Control, Electronics and Computer Science, Silesian University of Technology, Akademicka 16, 44-100 Gliwice, Poland

2. Department of Data Sciences, Dana-Farber Cancer Institute, Boston, MA, USA

3. Department of Biomedical Informatics, Harvard Medical School, Boston, MA, USA

#### 1 Reference-based compression

| Data set | CRAM 3.1 | RENANO | | CoLoRd-ref | | CoLoRd | |
|---|---|---|---|---|---|---|---|
| | Rel. size [%] | Rel. size [%] | DNA ratio [bpb] | Rel. size [%] | DNA ratio [bpb] | Rel. size [%] | DNA ratio [bpb] |
| Sorghum | 19.5 | 15.9 | 1.31 | 15.1 | 1.20 | 17.2 | 1.55 |
| NA12878 | 16.4 | 14.6 | 0.99 | 13.8 | 0.90 | 15.7 | 1.24 |
| CHM13 Guppy | 12.2 | - | - | 10.7 | 0.60 | 11.3 | 0.71 |
| CHM13 Bonito | 3.1 | 11.7 | 1.87 | 2.8 | 0.43 | 3.6 | 0.56 |
| M.bovis Guppy | 15.9 | 12.6 | 0.86 | 10.9 | 0.62 | 11.2 | 0.67 |
| M.bovis Bonito | 6.5 | 11.5 | 1.82 | 3.8 | 0.58 | 4.1 | 0.64 |

Table 1: Comparison of reference-based algorithms (CRAM 3.1 and RENANO) with CoLoRd in the reference-based and non-reference variants. The table presents relative sizes of the entire archives in percentages as well as DNA compression ratios in bits per base. All algorithms were configured to embed the reference genome in the archive. CoLoRd was run with *memory* (default) priority. CRAM archives were produced with SAMtools 1.12. RENANO failed to process CHM13 Guppy data set.

### 2 Consensus and variant calling performance

(a) Racon consensus: CHM13 ONT

| Quality mode | Breakpoints | Bases cov. 1 contig | Substitutions | Insertions | Deletions |
|---|---|---|---|---|---|
| lossless | 2 | 3,053,312,982 | 178,753 | 1,495,652 | 990,410 |
| *4-avg* (ONT default) | 2 | 3,053,310,979 | 175,330 | 1,530,904 | 971,829 |
| *4-fixed* | 2 | 3,053,178,229 | 179,923 | 1,519,670 | 1,002,576 |
| *none* Q10 | 11 | 3,051,652,997 | 158,465 | 1,534,387 | 965,221 |

(b) Racon consensus: CHM13 HiFi

| Quality mode | Breakoints | Bases cov. 1 contig | Substitutions | Insertions | Deletions |
|---|---|---|---|---|---|
| lossless | 0 | 3,054,797,046 | 3,044 | 2,961 | 31,427 |
| *5-avg* (HiFi default) | 0 | 3,054,797,823 | 3,016 | 2,996 | 28,730 |
| *5-fixed* | 0 | 3,054,794,313 | 3,179 | 3,253 | 34,573 |
| *none* Q10 | 0 | 3,054,797,221 | 3,185 | 2,264 | 26,973 |

(c) DeepVariant variant calling: HG002 HiFi

| Quality | SNP | | | indel | | |
|---|---|---|---|---|---|---|
| mode | prec. | recall | F1 | prec. | recall | F1 |
| lossless | 99.92 | 99.87 | 99.90 | 98.95 | 98.75 | 98.85 |
| *5-avg* (HiFi default) | 99.92 | 99.87 | 99.89 | 98.87 | 98.85 | 98.86 |
| *5-fixed* | 99.92 | 99.87 | 99.90 | 98.90 | 98.94 | 98.92 |
| *none* Q40 | 99.88 | 99.85 | 99.87 | 91.09 | 96.74 | 93.83 |
| *none* Q10 | 43.81 | 38.73 | 41.12 | 38.57 | 38.54 | 38.56 |

Table 2: Effect of lossy quality compression on downstream analyzes. **a-b** Results of the Racon consensus generation expressed by the number of substitutions, insertions, and deletions w.r.t. reference genome. **c** Accuracy of variant calling with DeepVariant.

### 3 Pipelines

#### Compression/decompression

```
# pigz
pigz -k -9 -p 24 <input>
pigz -d -k <archive>

# 7zip
7z a -mmt24 -mx9 <archive> <input>
7z e -mmt24 <archive>

# ENANO
enano -t 24 <input> <archive>
enano -d -t 24 <archive> <output>

# SPRING
spring -c -i <input> -o <archive> -l -t 24
spring -d -i <archive> -o <output> -t 24

# CoLoRd - lossy
colord compress-ont   <input> <archive> -t 24  # ONT
colord compress-pbhifi   <input> <archive> -t 24 # Hifi
colord compress-pbraw    <input> <archive> -t 24 # CLR
colord decompress <archive> <output>

# CoLoRd - lossless
colord compress-ont  -q org <input> <archive> -t 24  # ONT
colord compress-pbhifi  -q org <input> <archive> -t 24 # Hifi
colord decompress <archive> <output>

# RENANO
minimap2 -x map-ont --secondary=no --cs -t 24 <ref> <input> > <paf>
renano -t 24 -s <ref><paf> <input> <archive>

# CRAM
minimap2 -ax map-ont --secondary=no -t 24 <ref> <input> > <sam>
samtools view -T <ref> -@24 -O cram,version=3.1,small <sam> -o <archive>

# CoLoRd-ref
colord compress-ont -G <ref> -s <input> <archive> -t 24
```

#### Consensus—ONT

```
# recompress reads with lossy quality
./colord compress-ont -q <quality_mode> CHM13-ONT CHM13-ONT.fastq.colord
./colord decompress CHM13-ONT.colord CHM13-ONT.lossy.fastq

# map recompressed reads against reference genome - minimap 2.20
minimap2 -ax map-ont -t64 --secondary=no chm13.draft_v1.1.fasta CHM13-ONT.lossy.fastq > CHM13-ONT.lossy.sam

# run racon 1.4.20
racon -t 64 CHM13-ONT.lossy.fastq CHM13-ONT-lossy.sam chm13.draft_v1.1.fasta > CHM13-ONT.consensus.fasta 2> racon.log

# evaluate using minimap 2.20
minimap2 -t 32 --paf-no-hit -cxasm20 -r2k -z1000,500 -K 4000M chm13.draft_v1.1.fasta CHM13-ONT.consensus.fasta 1>mm1.log 2>mm2.log
wc mm1.log >stat1.log 2>stat2.log
minimap2 -t 32 -cxasm5 --cs -K 4000M -r2k chm13.draft_v1.1.fasta CHM13-ONT.consensus.fasta 1>mm3.log 2>mm4.log
sort -k6,6 -k8,8n mm3.log | paftools.js call - >stat3.log 2>stat4.log
```

#### Consensus—PacBio HiFi

```
# recompress reads with lossy quality
./colord compress-pbhifi -q <quality_mode> CHM13-HiFi CHM13-HiFi.colord
./colord decompress CHM13-HiFi.colord CHM13-HiFi.lossy.fastq

# map recompressed reads against reference genome
minimap2 -ax asm20 -t64 --secondary=no chm13.draft_v1.1.fasta CHM13-HiFi.lossy.fastq > CHM13-HiFi.lossy.sam

# run racon 1.4.20
racon -t 64 CHM13-ONT.lossy.fastq CHM13-HiFi-lossy.sam chm13.draft_v1.1.fasta > CHM13-HiFi.consensus.fasta 2> racon.log

# evaluate using minimap 2.20
minimap2 -t 32 --paf-no-hit -cxasm20 -r2k -z1000,500 -K 4000M chm13.draft_v1.1.fasta CHM13-HiFi.consensus.fasta 1>mm1.log 2>mm2.log
wc mm1.log >stat1.log 2>stat2.log
minimap2 -t 32 -cxasm5 --cs -K 4000M -r2k chm13.draft_v1.1.fasta CHM13-HiFi.consensus.fasta 1>mm3.log 2>mm4.log
```

```
sort -k6,6 -k8,8n mm3.log | paftools.js call - >stat3.log 2>stat4.log
```

#### Variant calling

```
# recompress reads with lossy quality
./colord compress-pbhifi -q <quality_mode> HG002.fastq HG002.colord
./colord decompress HG002.colord HG002.lossy.fastq

# map recompressed reads against reference genome
minimap2 -ax asm20 --secondary=no -R '@RG\tID:HG002\tSM:HG002' -t 48 GRCh38_no_alt_analysis_set.fasta HG002.lossy.fastq > HG002.lossy.sam

# convert SAM to BAM
samtools view -@ 48 -b -h HG002.lossy.sam > HG002.lossy.bam

# sort BAM
samtools sort -T ./temp -@ 48 -O bam HG002.lossy.bam > HG002.lossy.sorted.bam

# index BAM
samtools index HG002.lossy.sorted.bam

#  DeepVariant 1st pass
sudo docker run \
    gcr.io/deepvariant-docker/deepvariant:"1.1.0" \
    /opt/deepvariant/bin/run_deepvariant \
    --model_type PACBIO \
    --ref GRCh38_no_alt_analysis_set.fasta \
    --reads HG002.lossy.sorted.bam \
    --output_vcf pass1.vcf.gz \
    --num_shards 48

# index VCF
tabix -p vcf pass1.vcf.gz

# phase SNPs
whatshap phase \
    --output  pass1.phased.vcf.gz \
    --reference GRCh38_no_alt_analysis_set.fasta \
    pass1.vcf.gz \
    HG002.lossy.sorted.bam

# haplotag BAM
whatshap haplotag \
    --output HG002.lossy.haplo.bam \
    --reference GRCh38_no_alt_analysis_set.fasta \
    pass1.phased.vcf.gz \
    HG002.lossy.sorted.bam

# index haplotagged BAM
samtools index HG002.lossy.haplo.bam

#  DeepVariant 2nd pass
sudo docker run \
    gcr.io/deepvariant-docker/deepvariant:"1.1.0" \
    /opt/deepvariant/bin/run_deepvariant \
    --model_type PACBIO \
    --ref GRCh38_no_alt_analysis_set.fasta \
    --reads HG002.lossy.haplo.bam \
    --use_hp_information \
    --output_vcf pass2.vcf.gz \
    --num_shards 48

# compare variants against GIAB
python2 happy.py \
    HG002_GRCh38_1_22_v4.2.1_benchmark.vcf.gz \
    pass2.vcf.gz \
    -f HG002_GRCh38_1_22_v4.2.1_benchmark_noinconsistent.bed \
    -o eval.pass2 \
    -r GRCh38_no_alt_analysis_set.fasta \
```
